# The Therapeutic Antibody Profiler (TAP): Five Computational Developability Guidelines

**DOI:** 10.1101/359141

**Authors:** Matthew I. J. Raybould, Claire Marks, Konrad Krawczyk, Bruck Taddese, Jaroslaw Nowak, Alan P. Lewis, Alexander Bujotzek, Jiye Shi, Charlotte M. Deane

**Affiliations:** Department of Statistics, University of Oxford, Oxford, OX1 3LB, UK; Department of Antibody Discovery and Protein Engineering, MedImmune, Milstein Building, Granta Park, Cambridge, CB21 6GH, UK; Computational and Modelling Sciences, GlaxoSmithKline Research and Development, Stevenage, SG1 2NY, UK; Roche Pharma Research and Early Development, Large Molecule Research, Roche Innovation Center Munich, Penzberg, Germany; Chemistry Department, UCB Pharma, Slough, SL1 3WE, UK

**Keywords:** therapeutic monoclonal antibodies, developability guidelines, immunoglobulin gene sequencing, surface hydrophobicity, surface charge

## Abstract

Therapeutic monoclonal antibodies (mAbs) must not only bind to their target but must also be free from 'developability issues', such as poor stability or high levels of aggregation. While small molecule drug discovery benefits from Lipinski's rule of five to guide the selection of molecules with appropriate biophysical properties, there is currently no *in silico* analog for antibody design. Here, we model the variable domain structures of a large set of post-Phase I clinical-stage antibody therapeutics (CSTs), and calculate an array of metrics to estimate their typical properties. In each case, we contextualize the CST distribution against a snapshot of the human antibody gene repertoire. We describe guideline values for five metrics thought to be implicated in poor developability: the total length of the Complementarity-Determining Regions (CDRs), the extent and magnitude of surface hydrophobicity, positive charge and negative charge in the CDRs, and asymmetry in the net heavy and light chain surface charges. The guideline cut-offs for each property were derived from the values seen in CSTs, and a flagging system is proposed to identify nonconforming candidates. On two mAb drug discovery sets, we were able to selectively highlight sequences with developability issues. We make available the Therapeutic Antibody Profiler (TAP), an open-source computational tool that builds downloadable homology models of variable domain sequences, tests them against our five developability guidelines, and reports potential sequence liabilities and canonical forms. TAP is freely available at http://opig.stats.ox.ac.uk/webapps/sabdab-sabpred/TAP.php.

---

Therapeutic monoclonal antibodies (mAbs) must not only bind to their target but must also be free from ‘developability issues’, such as poor stability or high levels of aggregation. While small molecule drug discovery benefits from Lipinski’s rule of five to guide the selection of molecules with appropriate biophysical properties, there is currently no *in silico* analog for antibody design. Here, we model the variable domain structures of a large set of post-Phase I clinical-stage antibody therapeutics (CSTs), and calculate an array of metrics to estimate their typical properties. In each case, we contextualize the CST distribution against a snapshot of the human antibody gene repertoire. We describe guideline values for five metrics thought to be implicated in poor developability: the total length of the Complementarity-Determining Regions (CDRs), the extent and magnitude of surface hydrophobicity, positive charge and negative charge in the CDRs, and asymmetry in the net heavy and light chain surface charges. The guideline cut-offs for each property were derived from the values seen in CSTs, and a flagging system is proposed to identify non-conforming candidates. On two mAb drug discovery sets, we were able to selectively highlight sequences with developability issues. We make available the Therapeutic Antibody Profiler (TAP), an open-source computational tool that builds downloadable homology models of variable domain sequences, tests them against our five developability guidelines, and reports potential sequence liabilities and canonical forms. TAP is freely available at *http://opig.stats.ox.ac.uk/webapps/sabdab-sabpred/TAP.php*

## Introduction

Monoclonal antibodies (mAbs) are increasingly used as therapeutics targeting a wide range of membrane-bound or soluble antigens - of the 73 antibody therapies approved by the EU or FDA since 1986 (valid as of June 12^th^, 2018), 10 were first approved in 2017 (1). There are many barriers to therapeutic mAb development, besides achieving the desired affinity to the antigen. These include intrinsic immunogenicity, chemical and conformational instability, self-association, high viscosity, polyspecificity, and poor expression. *In vitro* screening for these negative characteristics is now routine in industrial pipelines (2).

While some cases of poor developability are subtle in origin, others are less ambiguous. High levels of hydropho- bicity, particularly in the highly variable complementarity-determining regions (CDRs), have repeatedly been implicated in aggregation, viscosity and polyspecificity (2–8). Asymmetry in the net charge of the heavy and light chain variable domains is also correlated with self-association and viscosity at high concentrations (4, 9). Patches of positive (10) and negative (11) charge in the CDRs are linked to high rates of clearance and poor expression levels. Product heterogeneity (e.g. through oxidation, isomerisation, or glycosylation) often results from specific sequence motifs liable to post- or co-translational modification.

An improved understanding of the factors governing these biophysical properties has enabled the development of *in silico* assays, which are faster and cheaper than their experimental equivalents. Computational tools already facilitate the identification of sequence liabilities, such as sites of lysine glycation (12), aspartate isomerisation (13), asparagine deamidation (13), and the presence of cysteines or N-linked glycosylation sites (14). A primary focus in recent years has been in designing software that can better predict aggregation proclivity. Many algorithms designed for this purpose use only the antibody sequence (4, 7, 8), although some suggest an analogous equation to use if a structure is available (4). One purely structure-based method is the structural aggregation propensity (SAP) metric (5), later included in the Developability Index (6). This has been shown to detect aggregation-prone regions, such as surface patches (15), and to be able to rank candidates relative to a known antibody developability profile (11), using a closely related antibody crystal structure. It is likely that SAP’s atomic-resolution analysis would be too sensitive to use in comparing static homology models of diverse antibodies, given the current accuracy of structure prediction (16).

An alternative approach to predict antibodies likely to have poor developability profiles is to highlight those candidates whose characteristics differ greatly from clinically-tested therapeutic mAbs; a similar strategy in the field of pharmacokinetics led to the Lipinski rules for small molecule drug design (17). Here, we build three-dimensional models of a large set of post-Phase I therapeutics and survey their sequence and structural properties, all of which are contextualized against human immunoglobulin gene sequencing (Igseq) data.

Using the distributions of these properties, we build the Therapeutic Antibody Profiler (TAP), an open-source computational tool that highlights antibodies with anomalous values compared to therapeutics. TAP builds a downloadable structural model of an antibody variable domain sequence, and tests it against guideline thresholds of five calculated measures likely to be linked to poor developability. It also reports potential sequence liabilities and all non-CDRH3 loop canonical forms.

## Results

### Sequence Data

As a dataset of mAbs unlikely to suffer with developability issues, we used the variable domain heavy and light chain sequences of 137 clinical-stage antibody therapeutics (“137 CSTs”) (18). To contextualize the properties of the CST set, we used a snapshot of the human antibody chain repertoire from a recent Ig-seq study by UCB Pharma Ltd. (“human Ig-seq”). The human Igseq dataset was analyzed as a set of non-redundant heavy or light chains (“human Ig-seq non-redundant chains”), and as a set of non-redundant CDR sequences (“human Ig-seq non-redundant CDRs”). We chose this Ig-seq data as it contains simultaneously sequenced heavy and light chains, and so is a promising starting point for realistic *in silico* pairing, required to make complete structural models.

### Model Structures

High quality structural information is critical to accurately predict the surface properties of antibodies. As crystal structures are often unavailable, or difficult to attain, accurate modeling is a necessary step of an effective antibody profiler. Accordingly, all our comparisons are made between models (even when crystal structures are available) to avoid a bias in terms of structural quality. ABodyBuilder (19) was run on the 56 CSTs with a reference PDB (20) structure (as of May 4^th^, 2018). Sequence identical templates were not included, and each resulting model was aligned to its reference to evaluate the backbone root-mean-square deviation (RMSD) across all IMGT regions (see *SI Methods*). The mean framework and CDR RMSDs (Table S1) were commensurate with the current state of the art (16). For our structural property calculations, we class surface-exposed residues as having a side chain with relative accessible surface area (ASA_rel,X_) ≥7.5%, compared to Alanine-X-Alanine for each residue X (21, 22). Using this definition, we identified all exposed residues in the models and PDB structures. Of the 7,057 exposed crystal structure residues, only 265 (3.76%) were wrongly assigned as buried in the models.

The results suggest that ABodyBuilder models are accurate enough for our analysis, we used this software to model all 137 CSTs (“137 CST models”) and a diverse subset of paired human Ig-seq chains (19,019 “human Ig-seq models”, see *SI Methods*). We then performed a series of *in silico* assays to determine the TAP guideline metrics.

### IMGT CDR Lengths

Loop length has a major impact on the nature of antigen binding. For example, if an antibody has a long CDRH3 loop, it tends to form most of the interactions with an antigen, whilst shorter CDRH3 loops contribute to concave binding sites where other CDRs more often assist in binding (23).

The 137 CST and human Ig-seq sequences were IMGT-numbered (24), and IMGT CDR definitions were used to split the sequences by region. The 137 CST CDRH3 loops had a median length of 12, compared to 14 for the human Ig-seq datasets (Fig. 1*A*). In the case of CDRL3 the distributions were closer, with a median length of 9 for the 137 CSTs, compared to 10 for the human Ig-seq datasets (Fig. S1*E*). To test whether hybridomal development might account for these findings - as it is known that mouse antibodies tend to have shorter CDRH3 loops than human antibodies (25) -we split the 137 CST dataset by developmental origin (Fig. S2). Fully human therapeutics were disproportionately represented at longer CDRH3s (mean: 13.21, median: 12), compared to chimeric, humanized, or fully murine therapeutics (mean: 11.91, median: 12). However, both therapeutic subsets still have shorter CDRH3s than human-expressed antibodies.

**Fig. 1.**
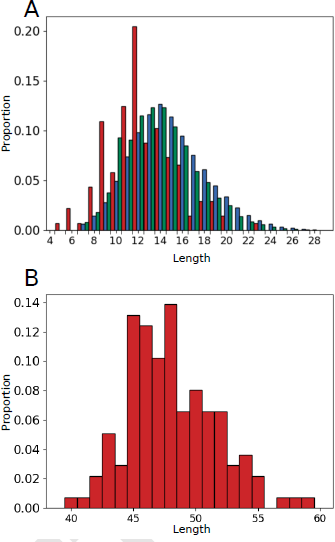
(*A*) Comparing the CDRH3 length distributions of the 137 CSTs (red), 1,696,918 human Ig-seq non-redundant CDRH3s (blue), and 4,587,907 human Igseq non-redundant heavy chains (green). The CSTs have a lower median CDRH3 length. (*B*) The distribution of total CDR length for the 137 CST dataset.

We also plotted the combined length of all CDRs for each antibody in the 137 CST dataset (Fig. 1*B*, median value 48). The human Ig-seq data is not paired, and so we could not plot its total CDR length. Our artificially paired human Igseq models were built to share a similar total CDR length distribution to the CSTs (Fig. S3), so that CDR length would not bias comparisons in other metrics. The 137 CST total CDR length was highly correlated to CDRH3 length (Pear- son’s correlation coefficient of +0.77, with a two-tailed p-value of 2.44e^-28^). As the total length of the CDRs captures both binding site shape (lower value, more concave) and CDRH3 length (typically shorter in CSTs than our human Igseq heavy chains), this metric was selected for inclusion in the final five TAP guidelines.

### Canonical Forms

In natural antibodies, all CDR loops, apart from CDRH3, are thought to fall into structural classes known as canonical forms (26, 27). We assigned length-independent canonical forms (see *Methods*) to the 137 CST and human Ig-seq models (Fig. S4). All assignable CST model CDRs were labeled with a canonical form also present in the human Ig-seq model dataset. Fewer than 19% of CST CDRs remained unassigned in each loop region, suggesting that, despite engineering, a clear majority of non-CDRH3 therapeutic CDR loops adopt well-characterized canonical forms. TAP reports the canonical form of each modeled loop, highlighting if any cannot be assigned.

### Hydrophobicity

Hydrophobicity in the CDR regions has been repeatedly linked to aggregation propensity in mAbs (2, 6–8). Using our homology models, we estimated the effective hydrophobicity of each residue by considering not only its degree of apolarity, but also whether or not it is solvent-exposed (side chain relative ASA_rel_ > 7.5% (21, 22)). As the energy of the hydrophobic effect is approximately proportional to the interface area (28), we developed a metric (Patches of Surface Hydrophobicity, PSH, see *Methods*) that yields higher scores if hydrophobic residues tend to neighbor one another in a region, rather than being evenly separated. We evaluated PSH for the 137 CST and human Ig-seq models across two regions (the CDR vicinity (see *Methods*) and the entire variable (Fv) region), and with five different hydrophobicity scales (29–33).

The results of all hydrophobicity scales were highly correlated (e.g. R^2^ ≥0.91 between all scales in the CDR vicinity). The mean CDR vicinity PSH values for the CST and Ig-seq distributions were 123.30 *±* 16.60 and 130.10 *±* 19.53 respectively (Kyte hydrophobicity scale (29), Fig. 2*A*). CSTs were noticeably underrepresented at higher CDR PSH values; galiximab is a rare example of a therapeutic antibody with a high value (Fig. 2*B*). The same divergence occurred to a much lesser extent across the entire Fv region, with mean values of 357.69 *±* 22.95 and 363.13 *±* 20.64 respectively (Fig. S5). This supports the theory that the high concentration conditions under which therapeutics are stored may render them less tolerant of large patches of hydrophobicity in the highly-exposed CDR vicinity, and also suggests that a subset of natural human antibodies would be unsuitable therapeutic candidates. We therefore included the CDR vicinity PSH score as a TAP guideline metric.

**Fig. 2.**
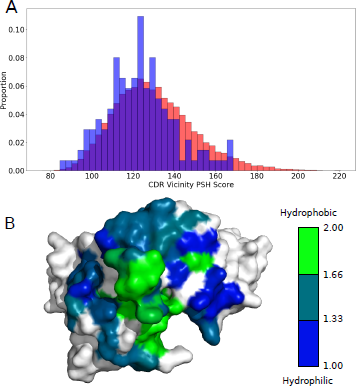
(*A*) CDR vicinity PSH scores across the 137 CST (blue) and human Ig-seq (red) models. The CSTs are underrepresented at higher PSH values. (*B*) Galiximab (Kyte & Doolittle CDR vicinity PSH score of 167.89) has a large surface-exposed patch of hydrophobicity in its CDRH3 loop. Heavy and light chain surfaces outside the CDR vicinity are colored in white.

### Charge

Surface patches of positive or negative charge have also been linked to negative biophysical characteristics (10, 11). We calculated two metrics designed to highlight regions of dense charge: the Patches of Positive Charge (PPC) and Patches of Negative Charge (PNC) measures (see *Methods*).

All surface residues were initially assigned the appropriate charge for their averaged pk_a_ values, as neighboring residues appear to have a limited effect at pH 7.4 (4). The charge of residues found to be engaging in salt bridges was then revised to 0.

The 137 CSTs tend to avoid patches of charge in their CDR vicinities, with 88.32% and 80.30% of them having PPC (Fig. 3*A*) and PNC (Fig. 3*B*) values below 1, respectively. Human Ig-seq models displayed similar PPC and PNC distributions, with the top 5% of PPC values being noticeably smaller than the top 5% of PNC values. Both PPC and PNC assays were carried forward as TAP guideline metrics.

**Fig. 3.**
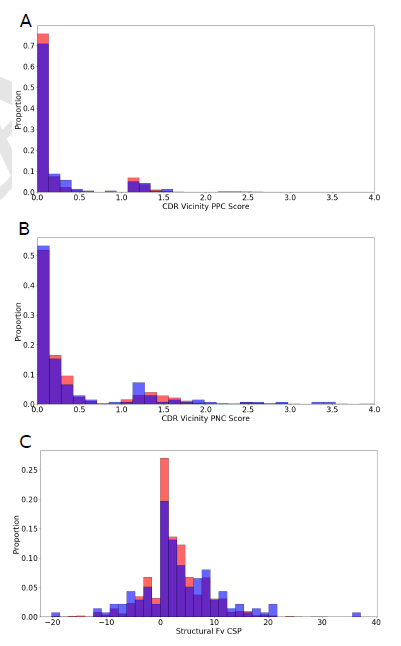
Histograms of 137 CST (blue) and human Ig-seq model (red) values of (*A*) patch-weighted positive charge (PPC) in the CDR vicinity, and (*B*) patch-weighed negative charge (PNC) in the CDR vicinity. In both metrics, the datasets are biased away from higher scores. (*C*) Histogram of Structural Fv Charge Symmetry Parameter values. Both datasets show a bias away from negative values.

When mAbs have oppositely charged V_H_ and V_L_ chains, they typically have higher *in vitro* viscosity values (4). This aggregate-inducing electrostatic attraction is captured at the sequence level by the Fv Charge Symmetry Parameter (FvCSP) metric - the mAb tends to be more viscous if the product of net V_H_ and V_L_ charges is negative (4). Harnessing our structural models, we calculated a variant (the Structural Fv Charge Symmetry Parameter, SFvCSP), which only includes residues that are surface-exposed, and not locked in salt bridges, in the evaluation of net charge. In galiximab, for example, we ‘correct’ the charge of arginine H108 and aspartic acid L56 to 0, as the model indicates that they form a salt bridge. The charges of the glutamic acid at position H6, the aspartic acids at positions H107, L98, and L108, and the histidine at position L40 are ignored as their side chains are buried. The FvCSP score for this antibody would be 0 (net heavy chain charge of 0, net light chain charge of −2.9), whilst the SFvCSP score is +2.0 (net heavy chain charge of +2, net light chain charge of +1). A similarly low percentage of CST models (21.9%) and human Ig-seq models (19.7%) had negative SFvCSP scores (Fig. 4*C*), with mean values of 3.34 *±* 7.44 and 2.52 *±* 5.54 respectively. With such a bias away from negative products, we chose the SFvCSP as our final TAP guideline property.

**Fig. 4.**
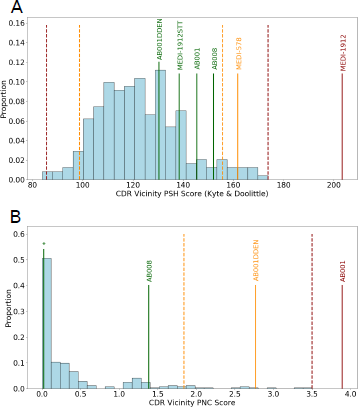
The (*A*) CDR vicinity PSH and (*B*) CDR vicinity PNC metrics for the combined set of 242 CSTs (light blue), and MedImmune case studies (colored by assigned flag). MEDI-578, MEDI-1912, and MEDI-1912SST all have the CDR vicinity PNC value labeled by an asterisk. Amber and red dashed lines delineate the 242 CST guideline thresholds. Case studies with prohibitive developability issues (MEDI-1912, AB001) are red-flagged for the PSH and PNC metrics respectively. Engineered versions without developability issues (MEDI-1912STT, AB001DDEN) return to the range of values previously seen in CSTs for all metrics.

### Developability Guidelines

While CSTs predictably share many features in common with human antibodies, our CDR length and hydrophobicity distributions imply that not every human antibody would make a good therapeutic. Consequently, our developability guidelines were set solely by CST values across the five selected metrics (Table 1). Amber flags indicate that the antibody lies within the extremes of the distributions, and should be interpreted as ‘be aware’. Red flags indicate a previously-unobserved value for that property, and should be interpreted as ‘cause for concern’.

**Table 1.** TAP amber and red flag cut-off thresholds, with respect to the clinical-stage therapeutic distributions. Len = Length.

| Metric | Amber Region | Red Region |
| --- | --- | --- |
| 1. Total CDR Len. | Bottom 5%/Top 5% | Above/Below |
| 2. PSH | Bottom 5%/Top 5% | Above/Below |
| 3. PPC | Top 5% | Above |
| 4. PNC | Top 5% | Above |
| 5. SFvCSP | Bottom 5% | Below |

To confirm that these threshold definitions do not typically flag mAbs without developability issues, we identified a further 105 whole antibody therapies (“105 CSTs”, listed in Dataset S1), not included in the 137 CST dataset, that had advanced to at least Phase II in clinical development.

Only eight of this set (7.69%) were assigned a red developability flag according to the boundaries set by the 137 CSTs, an average of 0.08 red flags per newly-tested therapeutic (Table S3). Erenumab received the most red flags - for total CDR length (60), CDR vicinity PSH (173.85), and CDR vicinity PPC (1.53). All other red-flagged therapeutics received only one: rafivirumab for total CDR length (60), intetumumab for CDR PSH (83.84), adacanumab, derlotuximab, lanadelumab and teprotumumab for CDR PPC (2.67, 2.66, 2.48, and 3.16 respectively), and quilizumab for Fv charge asymmetry (−20.40). The low red-flagging rate confirms that these guideline characteristics are highly conserved across therapeutic-like antibodies. Incorporating both sets of CSTs into a larger dataset (“242 CSTs”) led to the new guideline values shown in Table 2. While this had a large effect on the PPC thresholds, all other metrics were only slightly adjusted.

**Fig. 5.**
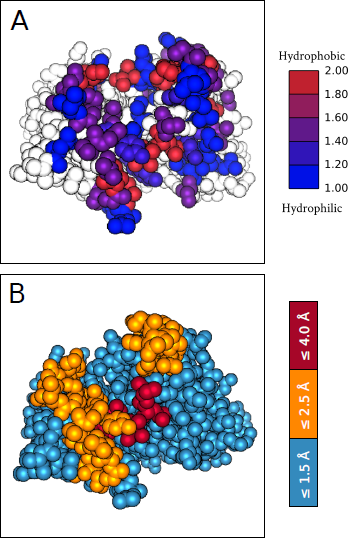
(A) An example TAP web application output showing the heavy atoms of an antibody as spheres colored by the hydrophobicity (Kyte & Doolittle scale, normalized between 1 and 2) of each residue in the CDR vicinity. (B) The ABody-Builder predicted model accuracy assignments (19) for each IMGT region, with heavy atoms shown as spheres. These are colored according to three backbone RMSD thresholds at a 75% confidence interval (both thresholds and confidence intervals can be modified in the web application). Better quality models will yield more reliable TAP metric values.

**Table 2.** TAP amber and red flag regions, as defined by the entire set of 242 CSTs. PSH score is calculated with the Kyte & Doolittle hydrophobicity scale (29). Metric numbers refer to Table 1. L = Length.

| Metric | Amber Region | Red Region |
| --- | --- | --- |
| 1. | $54 \leq L < 60$ | $L > 60$ |
| 2. | $83.84 \leq \text{PSH} < 100.71$<br>$156.20 < \text{PSH} \leq 173.85$ | $\text{PSH} < 83.84$<br>$\text{PSH} > 173.85$ |
| 3. | $1.25 \leq \text{PPC} < 3.16$ | $\text{PPC} > 3.16$ |
| 4. | $1.84 \leq \text{PNC} < 3.50$ | $\text{PNC} > 3.50$ |
| 5. | $-20.40 \leq \text{SFvCSP} < -6.30$ | $\text{SFvCSP} < -20.40$ |

### Case Studies

We tested whether these updated guideline values could highlight candidates with developability problems by running TAP on two datasets supplied by MedImmune (Fig. 4). A lead anti-NGF antibody, MEDI-578, showed minor aggregation issues during *in vitro* testing, of a level usually rectifiable in development, whereas the affinity matured version, MEDI-1912, exhibited unrectifiably high levels of aggregation (34). This observation was rationalized through SAP score (6) values, indicating that a large hydrophobic patch on the surface was responsible. TAP assigns MEDI-578 an amber flag, and MEDI-1912 a red flag - by a large margin - in the CDR vicinity PSH metric (Fig. 4*A*). Back-mutation of three hydrophobic residues in MEDI-1912 to those of MEDI-578 led to MEDI-1912STT, fixing the aggregation issue while maintaining potency. TAP assigns MEDI-1912STT no developability flags (Fig. 4*A*). A lead anti-IL13 candidate, AB008, had no developability issues, but the affinity-matured version, AB001, had very poor levels of expression (seven times lower than AB008) (11). The authors highlighted the role of four consecutive negatively charged residues in the L2 loop – mutation of the fourth negatively charged residue to neutral asparagine (AB001DDEN) was able to stabilize the loop backbone, mitigating the ionic repulsion of the DDE motif, and returning acceptable levels of expression. TAP assigns no developability flags to AB008, but a red flag to AB001, and an amber flag to AB001DDEN for its CDR vicinity PNC metric (Fig. 4*B*), again red-flagging the candidate with prohibitive developability issues. Both AB001 and AB008, confirmed monomers in solution (11), did not flag for CDR vicinity PSH score (Fig. 4*A*).

### Web Application

We have packaged the Therapeutic Antibody Profiler (TAP) into a web application, available at *http://opig.stats.ox.ac.uk/webapps/sabdab-sabpred/TAP.php*. TAP only requires the heavy and light chain variable domain sequences as an input, returning a detailed profile of your antibody with a typical runtime of less than 30 seconds. Flags (green, amber, or red) are assigned to each of the TAP metrics, with accompanying histograms. An interactive molecular viewer allows the user to visualize hydrophobicity (Fig. 5*A*), charge, and probable sequence liabilities on the antibody model surface. Estimated model quality can be easily accessed to help guide interpretation of the results (Fig. 5*B*). Finally, canonical forms are assigned to each non-CDRH3 loop. A full sample output is shown in Fig. S6.

## Discussion

By analyzing characteristics linked to poor developability, we have found evidence that suggests that not every human antibody would make a good therapeutic. This would be somewhat intuitive, as therapeutics suffer a range of stresses during development (including variation in pH and temperature, sheer forces, and high concentration storage conditions) that human-expressed antibodies are not exposed to. The TAP metrics therefore depend on the values seen across CSTs alone.

Our simple TAP guidelines will not capture the whole spectrum of developability issues. For example, they will not detect sources of immunogenicity, nor more subtle mechanisms that lead to poor stability. Nevertheless, we have shown that the TAP guidelines can selectively highlight antibodies with expression or aggregation issues (11, 34). The choice of 5^th^/95^th^ percentile amber flag cut-offs proved useful in distinguishing weak aggregators from non-aggregators (Fig. 4*A*). We intend to recalculate the threshold values on a monthly basis. This will take into account new mAbs that enter Phase II of clinical trials. It will also allow for the inevitable fluctuation in PSH, PPC, PNC, and SFvCSP values returned by every CST, as ABodyBuilder models will improve as the number of antibodies in the PDB increases (35). When sufficient numbers of mAbs have reached the market, we could increase the reliability of the TAP thresholds by only considering FDA/EU approved therapies, rather than all mAbs that have reached Phase II.

The thresholds themselves should not be interpreted as hard-and-fast rules, and the distance of red-flagged candidates outside the previously-observed bounds should be taken into consideration. Advances in process development and formulation may soon redefine the limits of permissible values (18).

## Methods

All modeling procedures are described in *SI Methods*.

### Human Ig-seq Data

Procured by UCB Pharma, this dataset contains 4,587,907 non-redundant heavy and 7,120,100 non-redundant light chains. The number of non-redundant CDR sequences is as follows: 174,490 CDRH1s, 279,873 CDRH2s, 1,696,918 CDRH3s, 455,125 CDRL1s, 8,708 CDRL2s, and 980,158 CDRL3s. For further information, including the chain pairing protocol, see *SI Methods*.

### Clinical-Stage Therapeutics

The initial set of 137 clinical-stage therapeutic (CST) antibody sequences were sourced from the Supporting Information of Jain *et al*. (18). The test set of 105 CST sequences was found through an extensive search of online resources, including the IMGT mAb [https://www.imgt.org/mAb-DB/] and Antibody Society [https://www.antibodysociety.org/late-stage-clinical-pipeline/] databases. The names and sequences of each CST are supplied in Dataset S1, with PDB structures (where available) listed at *http://opig.stats.ox.ac.uk/webapps/sabdabsabpred/Therapeutic.html*. All 242 CST ABody-Builder models are available on our group website, at *http://opig.stats.ox.ac.uk/webapps/sabdab-sabpred/TAP.php*.

### Canonical Forms

A length-independent canonical form clustering protocol (36) was run on the North-defined (37) CDR loops of a SAbDab (35) snapshot from 26th September 2017. Model loops were inferred to have identical canonical forms to the template used by ABodyBuilder (19).

### Surface-Exposed Residues

Residues are classed as surface-exposed if they have > 7.5% relative exposure (22) across side chain atoms, compared to the open-chain form Alanine-R-Alanine, as calculated with the Shrake & Rupley algorithm (21).

### CDR Vicinity

The “CDR vicinity” comprises every surface-exposed IMGT-defined CDR and anchor residue, and all other surface-exposed residues with a heavy atom within a 4 Å radius.

### Salt Bridges

Salt bridges were defined as pairs of lysines/arginines and aspartic acids/glutamic acids with a N^+-^O^-^ distance ≤3.2 Å.

### Hydrophobicity

Where R_1_ and R_2_ are two surface-exposed residues with a closest heavy atom distance, r_12_, < 7.5 Å and H(R,S) is the normalized hydrophobicity score (between 1 and 2) for residue R in scheme S, the Patches of Surface Hydrophobicity (PSH) metric can be calculated as:

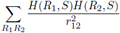

The hydrophobicity scales tested were: Kyte & Doolittle (29), Wimley & White (30), Hessa *et al*. (31), Eisenberg & McLachlan (32) and Black & Mould (33). Salt bridge residues were assigned the same value as glycine in each hydrophobicity scale.

### Charge

The following charges were assigned by sequence: Aspartic acid: −1, Glutamic acid: −1, Lysine: +1, Arginine: +1, Histidine: +0.1 (Henderson-Hasselbalch equation applied: pk_a_ 6, pH 7.4, and rounded-up to one decimal place). Tyrosine hydroxyl deprotonation was not considered. Salt bridge residues were assigned a charge of 0. The Patches of Positive Charge (PPC) and Patches of Negative Charge (PNC) metrics are analogous in form to PSH, with H(R,S) substituted for |Q(R)|, the absolute value of the charge assigned to residue R. Structural Fv Charge Symmetry Parameter (SFvCSP) values were calculated as:

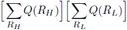

where R_H_, R_L_ are surface-exposed V_H_, V_L_ residues respectively.

## Acknowledgements

This work was supported by the Engineering and Physical Sciences Research Council (EPSRC) and the Medical Research Council (MRC) [grant number EP/L016044/1], GlaxoSmithKline plc, MedImmune Limited, F. Hoffmann-La Roche AG, and UCB Celltech. We thank Sebastian Kelm and James Heads for their helpful comments concerning our metrics. MIJR is grateful to Jinwoo Leem for his assistance in designing and implementing the web application.

